# LncDLSM: Identification of Long Non-coding RNAs with Deep Learning-based Sequence Model

**DOI:** 10.1101/2022.09.02.506180

**Authors:** Ying Wang, Pengfei Zhao, Hongkai Du, Yingxin Cao, Qinke Peng, Laiyi Fu

## Abstract

Long non-coding RNAs (LncRNAs) serve a vital role in regulating gene expressions and other biological processes. Differentiation of lncRNAs from protein-coding transcripts helps researchers dig into the mechanism of lncRNA formation and its downstream regulations related to various diseases. Previous works have been proposed to identify lncRNAs, including traditional bio-sequencing and machine learning approaches. Considering the tedious work of biological characteristic-based feature extraction procedures and inevitable artifacts during bio-sequencing processes, those lncRNA detection methods are not always satisfactory. Hence, in this work, we presented lncDLSM, a deep learning-based framework differentiating lncRNA from other protein-coding transcripts without dependencies on prior biological knowledge. lncDLSM is a helpful tool for identifying lncRNAs compared with other biological feature-based machine learning methods and can be applied to other species by transfer learning achieving satisfactory results. Further experiments showed that different species display distinct boundaries among distributions corresponding to the homology and the specificity among species, respectively. An online web server is provided to the community for easy use and efficient identification of lncRNA, available at http://39.106.16.168/lncDLSM.

## I. Introduction

Long non-coding RNAs (lncRNAs)—a type of ncRNAs (non-coding RNA) with the length of greater than 200 nucleotides—were initially thought to be spurious transcriptional noise due to their poor conservation. However, with the rapid growth of high-throughput sequencing technology, more and more researchers have confirmed that lncRNAs regulate gene expression in various stages and engage in different disease processes (Furlan & Rougeulle, 2016). For instance, studies have shown that AK023948 serves as a positive regulator for AKT (protein kinase B) (Mercer, Dinger, & Mattick, 2009), and the studies demonstrated that AK023948 upregulates in breast cancer. Therefore, it’s essential to identify lncRNAs to better understand the mechanisms of gene regulations and the inner causes of essential diseases.

The key of lncRNAs identification lies in separating lncRNAs and protein-coding transcripts (PCTs). However, lncRNAs have PCTs-like structures with a length of more than 200 nucleotides. After splicing, lncRNAs acquire 5’-methyl-guanosine caps (Wilusz, Sunwoo, & Spector, 2009) and poly-A tails (Rinn & Chang, 2012) structure like PCTs. In terms of transcription, both lncRNAs and PCTs are transcribed under the regulation of the common transcription factors, and the complex is mainly composed of RNA polymerase II (Pol II) (Y. J. Li, Egranov, Yang, & Lin, 2019). During differentiation, lncRNAs are dynamically expressed and spliced in different ways. Thus, many biological studies (Koirala, et al., 2017; Neil, et al., 2009) have shown that it is still a long way to identify those features to robustly discriminate lncRNA from these PCTs (David, et al., 2006).

To quantify the useful features to describe the lncRNAs, many lncRNA identification methods based on features extraction and machine learning/deep learning have been successively proposed (Achawanantakun, Chen, Sun, & Zhang, 2015; Baek, Lee, Kwon, & Yoon, 2018; Fan & Zhang, 2015; Ho Sung & B Franklin, 2012; Kang, et al., 2017; Kong, et al., 2007; Y. Li, et al., 2014; Pian, et al., 2016; Tuck & Tollervey, 2013; van Dijk, et al., 2011; Xu, et al., 2009), Among those feature extraction methods, the most common features are ORF-based features. Coding Potential Calculator (CPC) (Schneider, Raiol, Brigido, Walter, & Stadler, 2017) extracted sequence features containing the quality, coverage, and integrity of OR, and then used the Support Vector Machine (SVM) to assess the protein-coding potential of transcripts. Similarly, Coding Potential Assessment Tool (CPAT) (K. Sun, et al., 2013), lncRNA-MFDL (L. Sun, Liu, Zhang, & Meng, 2015), lncRNA-ID (L. Wang, et al., 2013) and lncRNApred (Kong, et al., 2007) all used the length and coverage of ORF as part of the features. In a recent study, lncRNAnet (L. Wang, et al., 2013) designed a CNN-based ORF indicator to automatically extract ORF-based features. But they still suffer from artificial feature calculation and lots of prior biological knowledge requirements for feature extraction, which is not always reliable. For example, because of the criterion that a transcript containing an ORF with the length above 100 amino acids is PCT (Fan & Zhang, 2015), the mouse Xist RNA gene with an ORF of length of 298 amino acids was initially mistaken for a PCT (Achawanantakun, et al., 2015). In addition, some researchers even proposed identification methods based on database searches and multiple sequence alignments (e.g. PhyloCSF) (Pian, et al., 2016). However, database searches still face the challenges of generalization issues for different conservative lncRNAs, poor annotation and long running time. Specially, most of the prior biological knowledge-based features (e.g. ORF-based features) are interrelated to the annotated sequence coding regions (CDs). The robustness of these methods is inevitably affected by the “silent” CDs of lncRNAs. Thus, all the sequence regions need to be considered when characterizing the transcripts. Partial region feature extraction strategy will inevitably cause biased long-range interaction of the transcripts, leading to unsatisfactory classification results. Lastly, differences between species also raised a significant challenge in differentiating between lncRNAs and mRNAs. For instance, 50% of both types of RNAs are between 200-900 bp range, whereas only about 25% were observed in this range from chickens, and those from cattle were in between(Kern, et al., 2019). Dealing with this issue during model design is also a task that cannot be neglected.

Based on this consideration, one of our primary tasks is extracting global features to characterize the transcripts without prior biological knowledge. Therefore, in this work, we mainly focus on two strategies for global feature extraction, i.e. k-mer frequency features and fast Fourier transform-based spectrum features. The differences of k-mer frequency features distribution reflect the differences among sequences composition regardless of specific biological significance. K-mer frequency features are similar to the bag of words (BOW) in natural language processing. BOW emphasizes the differences in the composition of words in a text but ignores the word order relations. In a financial report, the occurrence probability of “stock” is much higher than that of “basketball”. Similarly, some patterns of k-mer are more likely to appear in lncRNAs than PCTs and vice versa. Moreover, this phenomenon is more pronounced with the longer k-mer. And numbers of sequence identification studies have proved that k-mer frequency features are effective, although the k-mer frequency features ignore the order of nucleotides. On the other hand, based on the idea that gene transcription is a signal sampling process, transcripts can be regarded as a discrete time-domain signal, which can be processed by the fast Fourier transform (FFT) like other time-domain signals to obtain the spectrum features. It was demonstrated that the phenomenon of 3-based periodicity is widespread in the PCTs, which contain coding regions (Baek, et al., 2018; Okazaki, et al., 2002). Most of the Discrete Fourier Transform (DFT) based DNA/RNA sequence identification approaches compute the Fourier power spectrum features and obtain the signal-to-noise ratio information(Borsani, et al., 1991; Fickett, 1982; Fickett & Tung, 1992; M. F. Lin, Jungreis, & Kellis, 2011; Tiwari, Ramachandran, Bhattacharya, Bhattacharya, & Ramaswamy, 1997). Compared with k-mer frequency features, the spectrum features consider the order of nucleotides but ignore the sequence composition. Thus combining the two parts of features can comprehensively characterize the transcripts in terms of the feasibility of these two sets of features in the sequences identification task.

Benefiting from the ability to capture long-range interaction and association within biological sequences, deep learning techniques have recently facilitated advanced research in sequencing identification and classification. Thus, in our works, those two sets of long-range features are coupled with our designed two module based deep learning framework lncDLSM, to mine more potential patterns in the identification of lncRNAs. The whole lncDLSM consists of two parts, the first part is based on hierarchical input neural networks, called HINN-based analyzer, which is designed to extract the advanced features of the k-mer frequency features. Another part is a CNN-based detector, which is designed to extract the advanced features of the spectrum features. Then we merge these high-level features using another neural network-based prediction module to identify lncRNAs finally.

Finally, to tackle the problem of lncRNA identification across species. We designed a transfer learning strategy based on our framework to avoid massive parameter training procedures. Specifically, we trained the human model from scratch. We were then training other species models by fine-tuning the model. However, transfer learning requires similarity between the source and target domain. Otherwise, it is likely to cause negative transfer. Fortunately, species homology provides a guarantee for successful transfer. Although there is also species specificity, the transfer learning method can fine-tune the parameters of the human model to eliminate such effects. Specially, the results showed that transfer learning strategy may be an effective way to reveal species homology.

The main contributions of our work are summarized as follows:

1. Extract features based on the sequences only, without considering prior biological knowledge, to reduce the direct focus on coding regions. And lncDLSM achieved good performance with balanced sensitivity and specificity on five species data.
2. According to the traits of k-mer frequency features and spectrum features, HINN-based analyzer module and CNN-based detector module of lncDLSM are proposed to process the features compared to a single and simple solution in the past.
3. Transfer learning enhances the generalization of the lncDLSM and has the potential to become a novel method for the study of species homology.
4. An online web server is designed and provided for users to differentiate lncRNA and mRNA with a convenient access portal.

## II. Material and Methods

### A. Datasets

NONCODE (current version v5.0) is an integrated knowledge database designed for non-coding RNAs, especially lncRNAs (Anastassiou, 2000). The lncRNAs of 17 species can be downloaded from NONCODE (human, mouse, pig, cow, rat and zebrafish, etc.). In NONCODE database, the numbers of lncRNAs are 172216, 131697, 29585, 23515 and 24879 for human, mouse, pig, cow, and rat, respectively.

As shown in **Fig. 1**, the lncRNAs with a length above 3000 nucleotides cover only about 11.64% of the human data and approximately 10.30% of the mouse data. Only transcripts with a length less than 3000nt were used to train the model in our experiments. GENCODE (Pian, et al., 2016; Yin & Yau, 2007) is a open source annotation database with high quality data. The FASTA files of lncRNAs and PCTs of human and those of mouse can be downloaded. In our experiments, the PCTs of human (Release 30) and mouse (Release M21) were used. The total number of PCTs in human is 99731, and this of mouse is 66650. RefSeq (NCBI Reference Sequence Database) collection provides a comprehensive, non-redundant, annotated set of sequences, including DNA sequences, transcripts, and proteins (Anastassiou, 2000). From the RefSeq database, we downloaded PCTs of pig, cow and rat.

**Fig. 1.**
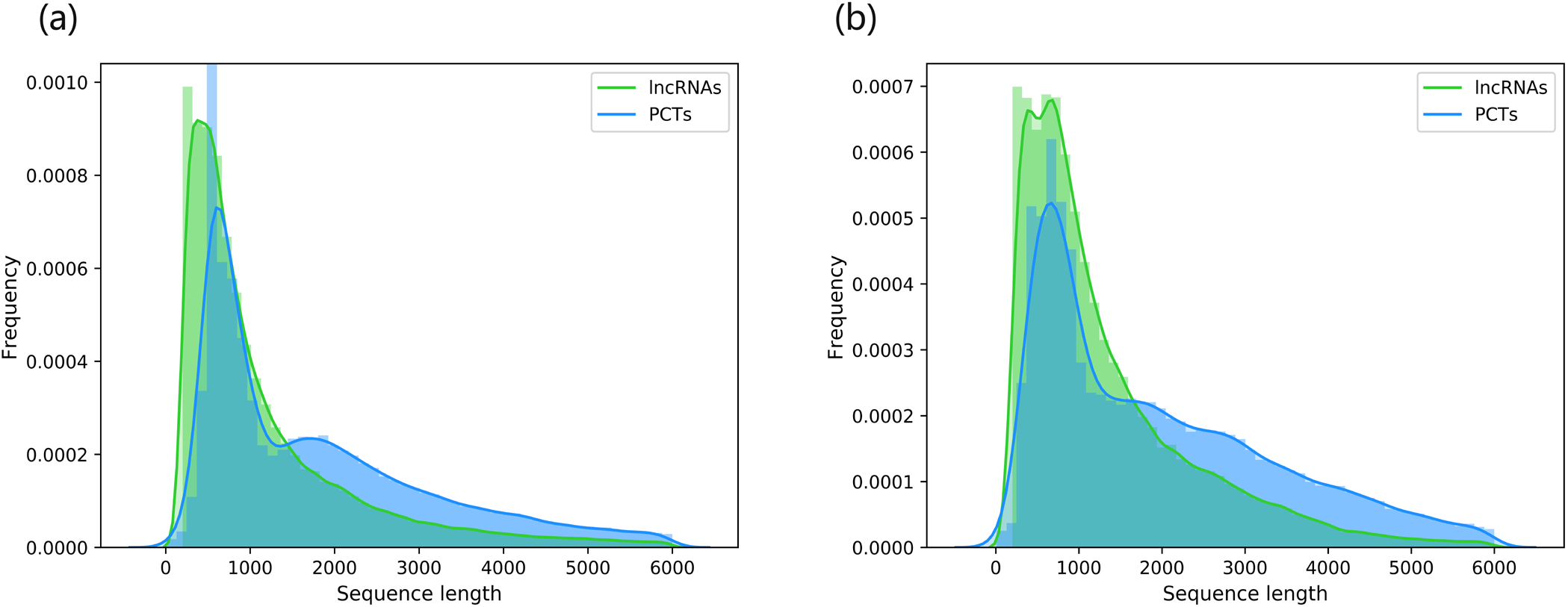
Length distribution of the PCTs and lncRNAs on the (a) human and (b) mouse data.

In our experiments, 70000 lncRNAs and 70000 PCTs of human (BMT) were used to train the model of human which is also the basic model to train the identification model of other species. After removing the training datasets from the databases, 3000 lncRNAs and 3000 PCTs, obtained from NONCODE and GENCODE respectively, of human were randomly selected to evaluate the model (HT). For diversity, we prepared lncRNAs of mouse, pig, cow and rat from NONCODE. PCTs of the mouse were obtained from GENCODE and those of the other three species from RefSeq. **TABLE I** lists the number of transcripts used in our work. In **TABLE I**, the testing datasets of mouse, pig, cow and rat were recorded as MT, PT, CT, and RT, respectively.

**TABLE I.** Units for Magnetic Properties

|  | BMT | HT | MT | PT | CT | RT |
| --- | --- | --- | --- | --- | --- | --- |
| LncRNAs | 70000 | 3000 | 3000 | 3000 | 3000 | 3000 |
| PCTs | 70000 | 3000 | 3000 | 3000 | 3000 | 3000 |
| Total | 140000 | 6000 | 6000 | 6000 | 6000 | 6000 |

### B. K-mer frequency features

A k-mer pattern is a simple sequence with length k, composed of nucleotides (A, G, C, T). Researchers in the field of biological sequencing believe that the species specificity of the k-mer frequency features becomes more apparent as the k value increases. Actually, the distributions of k-mer frequency features also differ in different types of sequences, and this difference will increase as the k value increases. However, the excessive pursuit of distribution differences can lead to the curse of dimensionality.

The analysis of k-mer frequency features with bigger k value usually requires deeper neural networks to capture more details. Therefore, HINN-based analyzer is designed to adapt to the discrepancy of the neural network depth of different dimensional data. In our study, 5-mer frequency features correspond to the deepest neural network, 4-mer frequency features second, and 3-mer frequency features least. Later experiments have also verified the effectiveness of this design.

### C. Fast Fourier Transform and 3-based periodicity

Fast Fourier transform is a fast implementation of Discrete Fourier Transform. It improves the DFT algorithm based on the odd, even, imaginary and real properties. It is a big step forward in applying the discrete Fourier transform to computer systems or digital systems and has been widely applied in the engineering field (Fang, et al., 2017; Harrow, et al., 2006; Jalali, Gandhi, & Scaria, 2016). In our study, the introduction of FFT solves the problem that binary sequences are too sparse. In addition, many studies have shown that CNN is a very suitable and effective neural network for processing spectrum features in speech recognition (O’Leary, et al., 2016; Rostami, Shanehsazzadeh, & Fardmanesh, 2018). Therefore, we abandoned the time-consuming RNNs in favor of convenient and efficient CNNs.

Although we did not directly extract the corresponding features based on prior biological knowledge, we cannot avoid 3-based periodicity when talking about FFT. The 3-based periodicity is described as a prevalent phenomenon that a pronounced peak at the frequency N/3 of the Fourier power spectrum curve of the DNA/RNA sequences with length N, which contains protein-coding regions. It was proved that the 3-based periodicity in a DNA/RNA sequence is mainly caused by the imbalance of nucleotides distribution in the coding region (Abdel-Hamid, Mohamed, Hui, & Penn, 2012; Deng, Abdel-Hamid, & Yu, 2013; Fickett, 1982; Peng, Liu, Tong, & Colavolpe, 2017; Yun, Han, Fan, Zhang, & Hong, 2018). The nucleotide distribution in the three codon positions is unbalanced in an exon region while the nucleotides distribute uniformly in the three codon positions in an intron region. The essence of this phenomenon is that the protein’s preference for special amino acid composition leads to the usage deviations of the nucleotides in the coding regions (Fickett, 1982; Fickett & Tung, 1992; Tiwari, et al., 1997; Yin & Yau, 2005, 2007). In our work, instead of considering the problem from this perspective, we fed the original spectrum features directly into the model. The CNN-based detector was used to mine more valuable information.

### D. Methods

The proposed method lncDLSM is shown in Figure 1. To determine the category of candidate transcripts, our method flow includes the following five phases: (a) extracting spectrum features, (b) extracting k-mer frequency features, (c) acquiring the advanced characterization from spectrum features by the CNN-based detector, (d) acquiring the advanced characterization from k-mer frequency features by the HINN-based analyzer, (e) features combination and prediction. The details of our method are given in Algorithm 1. Our model takes a set of sequences as the input and predicts these sequences as the output. The first phase (Lines 4-11) includes the separation of binarization, padding, FFT, and real and imaginary parts. In the step of extracting k-mer frequency features (Lines 12-13), k is 3, 4, and 5 respectively. Lines 17-23 describe the process of further characterization of features and the model’s training.

#### Algorithm 1 Pseudocode of IncDLSM

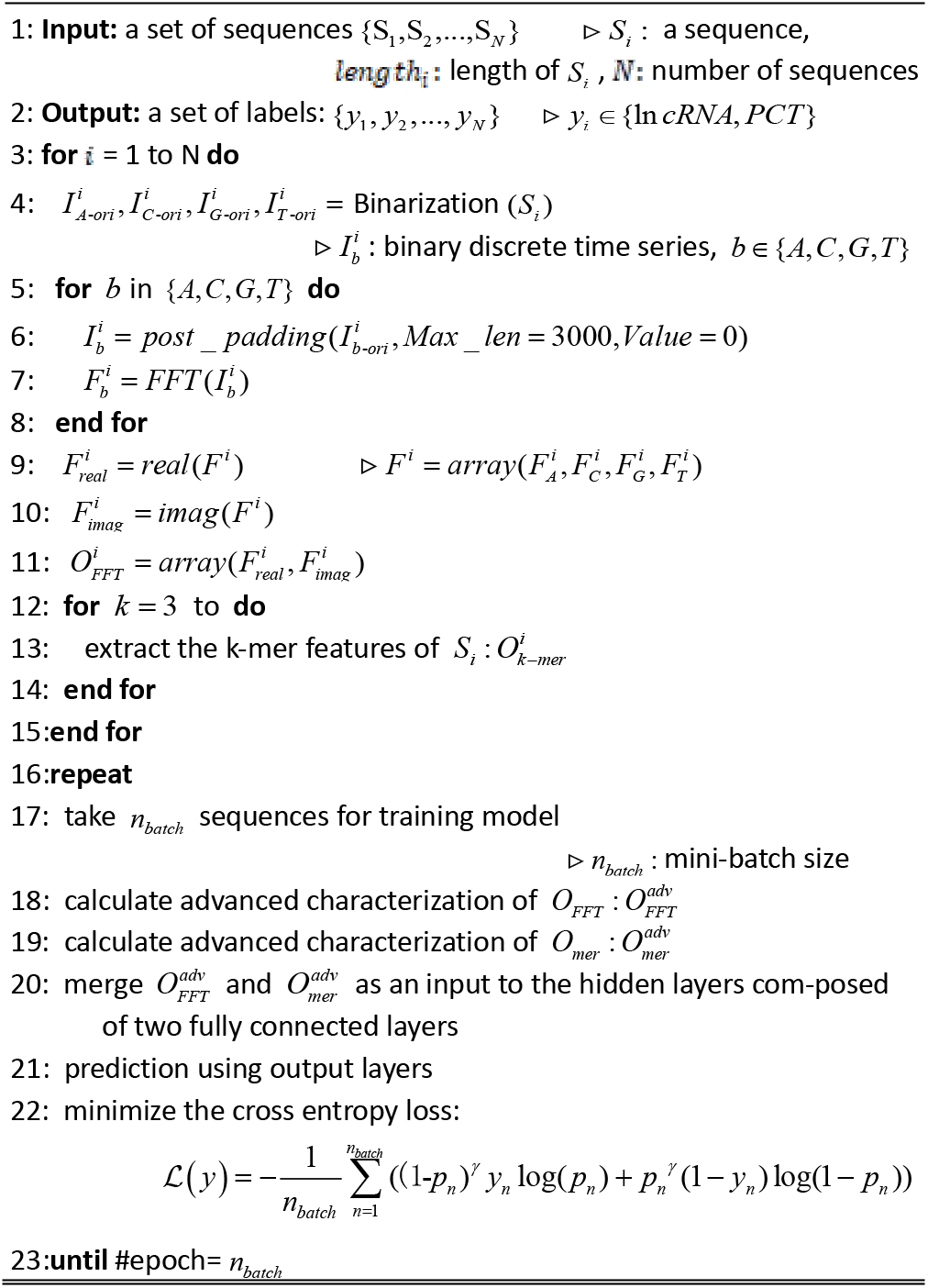

#### 1) CNN-based detector

Given a transcriptome sequence with length *L*. The first step is binarizing the sequence by (1).

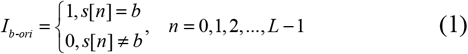

where *s*[*n*] represents a transcriptome sequence *I_b-ori_*, is a binary discrete time series, *b* ∈ *B* = {*A, C, G, T*}.

To facilitate the training of the neural network model, four discrete binary time series 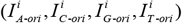 were padded to the same length with ‘0’. In our experiments, four new discrete discrete time series with length 3000 were obtained after the padding. Then, we performed FFT on each binary discrete time series, its representation in the frequency domain can be obtained. In this paper, the Discrete Fourier Transform formula is provided since FFT is just a fast implementation of DFT, and both have the same essence.

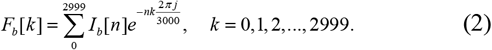

Since the spectrum is symmetrical, only the first half of the spectrum was used. In addition, instead of taking the spectrum directly as the features, the real and imaginary parts were taken as two channels. Thus the spectrum features of a transcriptome sequence with shape (4, 1500, 2) were obtained.

As mentioned above, CNNs are convenient and efficient neural networks for processing spectrum features. For the unique characteristics of spectrum features of the transcriptome sequence data, we designed a CNNs-based detector. Since the spectrum features have two channels, we first used a two-dimensional convolution to extract local features. The ‘A’, ‘C’, ‘G’ and ‘T’ ribonucleotides are equally important in the sequence. So in each convolution step, the local spectrum features of all ribonucleotides were extracted simultaneously. Thus, the size of convolution filters can meet the requirement that the first dimension is always equal to 4. In addition, many studies have shown that using multi-scale filters in one layer can lead to better model performance (Fickett & Tung, 1992; Szegedy, Ioffe, Vanhoucke, & Alemi, 2017; Tiwari, et al., 1997; Yin & Yau, 2005, 2007). Thus, we designed (4, 4), (4, 8) and (4, 16) three sizes filters with 16 per type. And the ‘padding’, a hyperparameter of Conv2D layers in the Keras, was set to ‘valid’. After two-dimensional convolution processing and dimensionality reduction by two-dimensional max-pooling layers, the three-dimensional feature map can generate to two-dimensional feature map. Next, feature extraction was further performed using one-dimensional convolutional layers and one-dimensional max-pooling layers. Finally, after several hidden layers processing, we got an 8-dimensional vector that can represent the variations of a transcriptome sequence 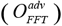.

#### 2) HINN-based analyzer

The experiment found that 3-mer, 4-mer and 5-mer frequency features achieved better recognition results than other patterns. In this paper, let *k* = 3,4,5. There are 4^3^ + 4^4^ + 4^5^ = 1344 patterns, and the step size is set to one nucleotide. For one sequence with length L, a total of 1344 k-mer frequency features can be computed by (3).

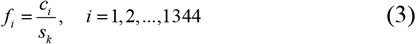

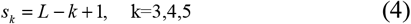

where *c_i_* is the number of k-mer patterns in a sequence, ^*f_i_*^ is one of the features. HINN-based analyzer was designed to specifically analyze and extract valid information of the k-mer frequency features. Different from the previous methods of feeding the whole k-mer frequency features to the neural network simultaneously, these features were divided into three groups: 3-mer frequency features with shape (64,1), 4-mer frequency features with shape (256,1) and 5-mer frequency features with shape (1024,1). It can be argued that as the value k increases, the patterns of the k-mer become more and more complex and contain more implicit information. Therefore, three groups of features need different numbers of hidden layers. In the HINN-based analyzer, 5-mer frequency features were processed by deepest neural network and 3-mer frequency features by shallowest neural networks. Firstly, feeding the 5-mer frequency features to the HINN-based analyzer, these features were processed by hidden layers and were processed to 256 dimensions. Secondly, merging the above 256-dimensional features and 4-mer frequency features, the obtained 512-dimensional features were compressed by hidden layers to get 64-dimensional features. Finally, merging the above 64-dimensional features and 3-mer frequency features, the obtained 128-dimensional features were mapped to an 8-dimensional vector after passing through the hidden layers. And the compositions of a transcriptome sequence were represented by this 8-dimensional vector 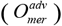.

#### 3) Learning lncRNAs

After processing the CNN-based detector and HINN-based analyzer, two 8-dimensional vectors 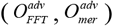 represent the variations and the compositions of a transcripts sequence. We merged as a 16-dimensional vector fed into the classifier composed of several fully connected layers. The final output of the classifier is a 2-dimensional vector modified as normalized probabilities for each class by the Sigmoid function. The Sigmoid function is often used as a threshold function for the neural networks, mapping variables between 0 and 1. The focal loss that adds a factor (1-*p_n_*)*^γ^* to the standard cross entropy criterion (Szegedy, et al., 2017) was selected as the loss function in the training periods. The parameter ^*γ*^ is a positive number. The addition factor forces the model to focus on the hard, misclassified samples. The focal loss is expressed as follows:

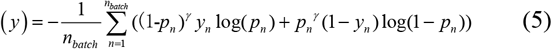

where *y_n_* is the true label, *p_n_* is the predicted probability, and *n_batch_* is the batch size.

#### 4) Transfer learning

Transfer learning is a special machine learning method that uses the model developed for task A as an initial point and reuses it in developing a model for task B. Compared to multi-task learning, transfer learning cares most about the target task rather than learning all the source and target tasks simultaneously. In transfer learning, the roles of the source and target tasks are no longer symmetric (Szegedy, Vanhoucke, Ioffe, Shlens, & Wojna, 2016). There are mainly instance-based transfer, features-based transfer, and shared parameters-based transfer. Shared parameters-based transfer is to research how to find common parameters or prior distribution between the models of the source v.s. target data, so that knowledge transfer can be further processed and achieved. It is assumed that each learning task-related model will share some of the same parameters or prior distributions. And fine-tuning, which was used in our study, is a very popular trick in shared parameters-based transfer.

In terms of application research, transfer learning has attracted the attention of bioinformatics researches. Tongxin Wang et al. effectively combined different batches of scRNA-seq data through transfer learning to remove batch effects and distinguish cell types (Xie, Girshick, Dollar, Tu, & He, 2017). Suyu Mei and Hao Zhu proposed AdaBoost-based multi-instance transfer learning for predicting proteome-wide interactions between salmonella and human proteins (Zoph, Vasudevan, Shlens, & Le, 2018).

#### 5) Model evaluation

To evaluate the performance of our model against other competitors. We selected multiple evaluation metrics, i.e. Sensitivity, Specificity, Accuracy, and F1 score, to conduct the comparison. Since all these selected metrics are frequently used for the evaluation on classification tasks. Therefore, we adopted them to calculate the results on all the compared methods. These metrics are defined as follows:

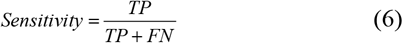

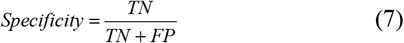

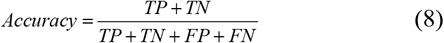

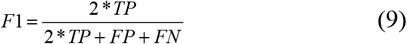

## III. Experimental results

As illustrated in the previous section, the existing methods use prior biological knowledge to identify lncRNAs. In our study, we extracted frequency features, and spectrum features, and these features have no biological significance by themselves. Thanks to deep neural networks’ powerful feature extraction capabilities, it is reliable to hand over the task of mining biological information hidden in our extracted features. Specially, lncDLSM will not suffer from deficiencies and defects in knowledge and will achieve good performance with balanced sensitivity and specificity. After obtaining a human model, the usage of transfer learning reduces the difficulty of model training for other species.

### A. Prediction of performance

The empirical optimal hyperparameters and model structures were established through various combinations of hyperparameters and massive contrast experiments. In this section, all the experiment results were obtained from the average performance based on the BMT’s five-fold cross-validation.

**Table II** shows the performance of the model for each component. For CNN-based detector (16 filters for each of the three sizes), we modify the sizes of the filter to (4, 4), (4, 8) and (4, 16) respectively, and the numbers of the filter to 8*3 and 32*3 respectively (x*3, the number of filters is x per type). As seen from the experimental results, too large or too small filter sizes got worse when using filters with one size. This observation can get an intuition explanation from two extreme cases: in one case, the size of the filter is (1, 1), and the model cannot explore the cross information among different features; in another case, that filter size is equal to the input resolution, a convolutional layer becomes a fully-connected layer, which has a disadvantage in accuracy and efficiency. As mentioned above, the convolutional layers with multi-scale filters gradually replace the convolutional layer with single-scale filters due to their excellent experimental performance. Compared to single-scale filters, CNN-based detector similarly acquired more satisfying results due to the usage of multi-scale filters. It is found that the number of convolutional filters equal to 16*3 reaches the highest accuracy. For HINN-based analyzer (using 3-mer, 4-mer, 5-mer; 5-mer corresponding to deepest neural networks), we performed these modifications: (1) inversing the order of k-mer (3-mer corresponding to deepest neural networks); (2) removing the hierarchical structure; (3) replacing the 3, 4, 5-mer with 2, 3, 4-mer and 4, 5, 6-mer respectively. Compared with Non-hierarchical HINN-based analyzer, both the unmodified and inversed HINN-based analyzer have obtained higher accuracy, confirming that hierarchical input is necessary. From the experimental results of unmodified and inversed HINN-based analyzer, it is proved that the design rationality of HINN-based analyzer is based on human cognitive processes. In the replacing features experiments, the 4, 5, and 6-mer frequency features acquired the best results with a slight advantage. Trading off the accuracy and efficiency, we chose 3, 4, and 5-mer frequency features to achieve the purpose of identification. LncDLSM, including CNN-based detector and HINN-based analyzer, achieved the best identification effect.

**Table II.**
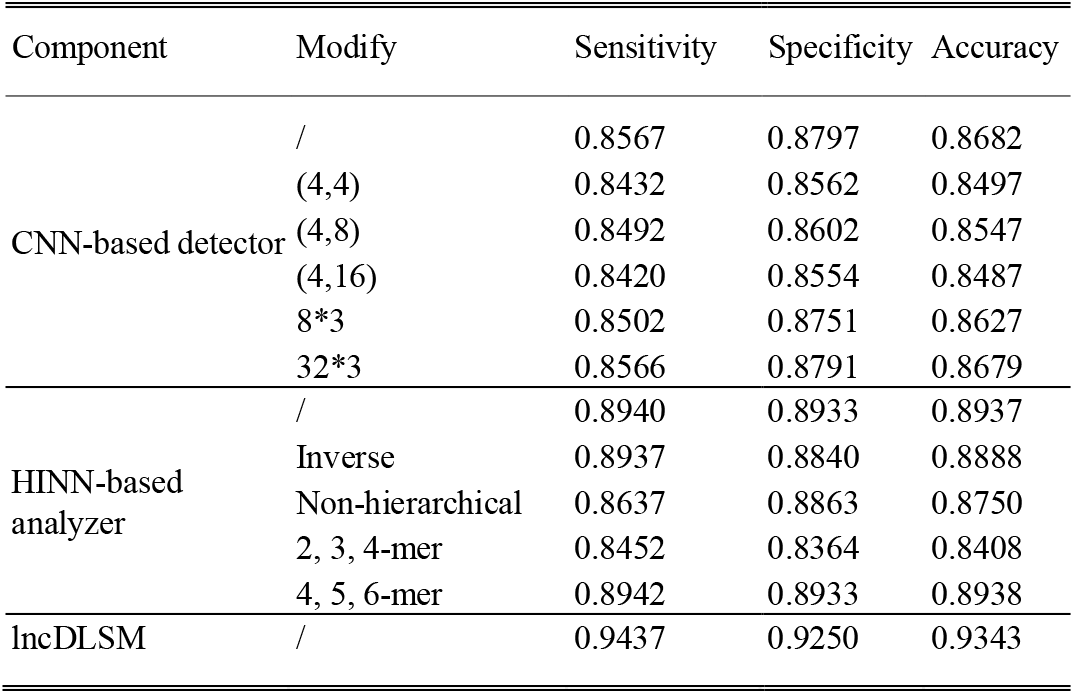
Effects of component modifications

**Table III** shows an obvious conclusion that focal loss is the optimal objective function for the model. Focal loss focuses on the hard, misclassified samples to promote the model parameters closer to the global solution. The ^*γ*^ of 1 showed the highest accuracy in experiments.

**Table III.** Effects of different loss functions

| Loss function | Sensitivity | Specificity | Accuracy |
| --- | --- | --- | --- |
| Focal loss ($\gamma=1$) | **0.9437** | **0.9250** | **0.9343** |
| Focal loss ($\gamma=0.5$) | 0.9310 | 0.9224 | 0.9267 |
| Focal loss ($\gamma=2$) | 0.9322 | 0.9216 | 0.9269 |
| Cross entropy loss | 0.9308 | 0.9205 | 0.9257 |
| Mean squared loss | 0.9299 | 0.9108 | 0.9204 |
| Mean absolute loss | 0.9203 | 0.9002 | 0.9103 |
| Hinge loss | 0.8901 | 0.8802 | 0.8852 |

### B. Comparison of prediction methods

In order to prove the validity of our model, lncDLSM was compared with the following recognition methods: lncRNAnet, a Neural Network-based model, uses RNNs to learn the intrinsic features of the sequences and proposes an ORF indicator to reinforce the model (T. Y. Lin, Goyal, Girshick, He, & Dollar, 2017); Coding-Potential Assessment Tool (CPAT) extracts ORF size, ORF coverage, Fickett TESTCODE statistic and hexamer usage bias four sequence features to build a logistic regression model (Pan & Yang, 2010); Coding Potential Calculator 2 (CPC2), an updated version of CPC (T. X. Wang, et al., 2019), uses a Support Vector Machines (SVM) model built with six biologically meaningful sequence features including log-odds score, ORF coverage, ORF integrity, hits number, hits score and frame score (Mei & Zhu, 2014); PLEK, an alignment-free tool based on an improved k-mer scheme, is also a SVM-based model with a radial basis functional kernel (Baek, et al., 2018). mRNA RNN (mRNN) uses Gated Recurrent Unit (GRU) RNNs to solve the “vanishing gradient problem” caused by long nucleic acid sequences (L. Wang, et al., 2013). RNAsamba, a Neural Network-based model. Starting from the initial nucleotide sequence, RNAsamba takes into account the information from two different sources, the entire nucleotide sequence and the longest ORF, to calculate the coding potential of a given transcript(Camargo, Sourkov, Pereira, & Carazzolle, 2020).

#### 1) Performance comparison of the human dataset

**Table IV** shows the identification performance on the testing dataset of human. LncDLSM achieved 93.43% prediction accuracy, which is 2.03%, 7.83%, 16.06%, 17.56%, 2.41%, and 2.50% higher than that of lncRNAnet, CPAT, CPC2, PLEK, mRNN and RNAsamba predictors, respectively. And lncDLSM gave the highest scores in terms of specificity and F1. Although CPAT had the highest sensitivity, it suffered from poor specificity, accuracy, and F1. CPC2 and PLEK had similar performance in all aspects. Furthermore, there was a phenomenon that the predictions of CPAT, CPC2, and PLEK were biased towards lncRNAs. LncRNAnet had relatively improved PCT prediction of PCTs but had not completely solved the problem. One possible explanation is that many lncRNAs have similar ORF to PCTs. LncRNAnet detected an ORF indicator by CNNs instead of computing the statistical properties of ORF, improving the identification ability of pseudo-ORF to some extent. In addition, lncRNAnet learned intrinsic features by RNNs, further enhancing the recognition ability of the model. PLEK ignores the variations of the sequence, resulting in the least specificity. LncDLSM, by contrast, achieved the best overall performance with balanced sensitivity and specificity. In addition, the highest F1 also proved this.

**Table IV.** Comparison of prediction performance (Species: Human)

| Method | Sensitivity | Specificity | Accuracy | F1 |
| --- | --- | --- | --- | --- |
| LncDLSM | 0.9437 | **0.9250** | **0.9343** | **0.9349** |
| LncRNAnet | 0.9307 | 0.8973 | 0.9140 | 0.9154 |
| CPAT | **0.9600** | 0.7520 | 0.8560 | 0.8696 |
| CPC2 | 0.9490 | 0.5983 | 0.7737 | 0.8074 |
| PLEK | 0.9447 | 0.5727 | 0.7587 | 0.7965 |
| mRNN | 0.9220 | 0.8983 | 0.9102 | 0.9112 |
| RNAsamba | 0.9140 | 0.9047 | 0.9093 | 0.9098 |

In **Fig. 3**, we drew the Receiver Operating Characteristic (ROC) curve and calculated the Area Under the Curve (AUC) corresponding to each method. Compared with lncRNAnet (0.9663), CPAT (0.9418), CPC2 (0.8794), PLEK (0.8801) mRNN (0.9582) and RNAsamba (0.9654), lncDLSM (0.9717) achieved the highest AUC score. It reinforces the above finding and indicates that lncDLSM is an excellent classifier.

**Fig. 2.**
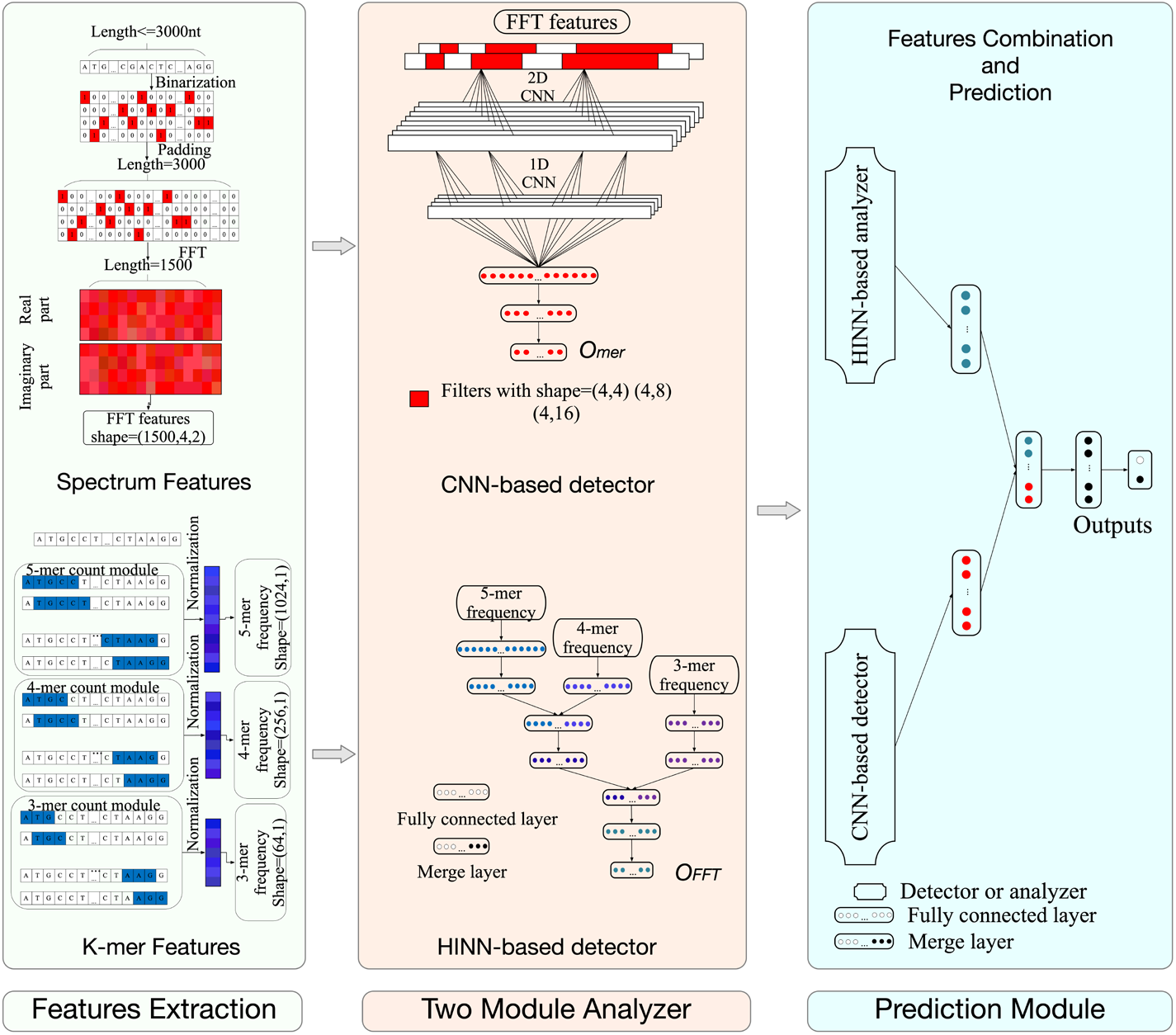
The overview of lncDLSM. (a) The scheme of the spectrum features extraction. (b) The scheme of the k-mer frequency features extraction. (c) CNN-based detector: a deep learning framework for extracting advanced features of spectrum features.

**Fig. 3.**
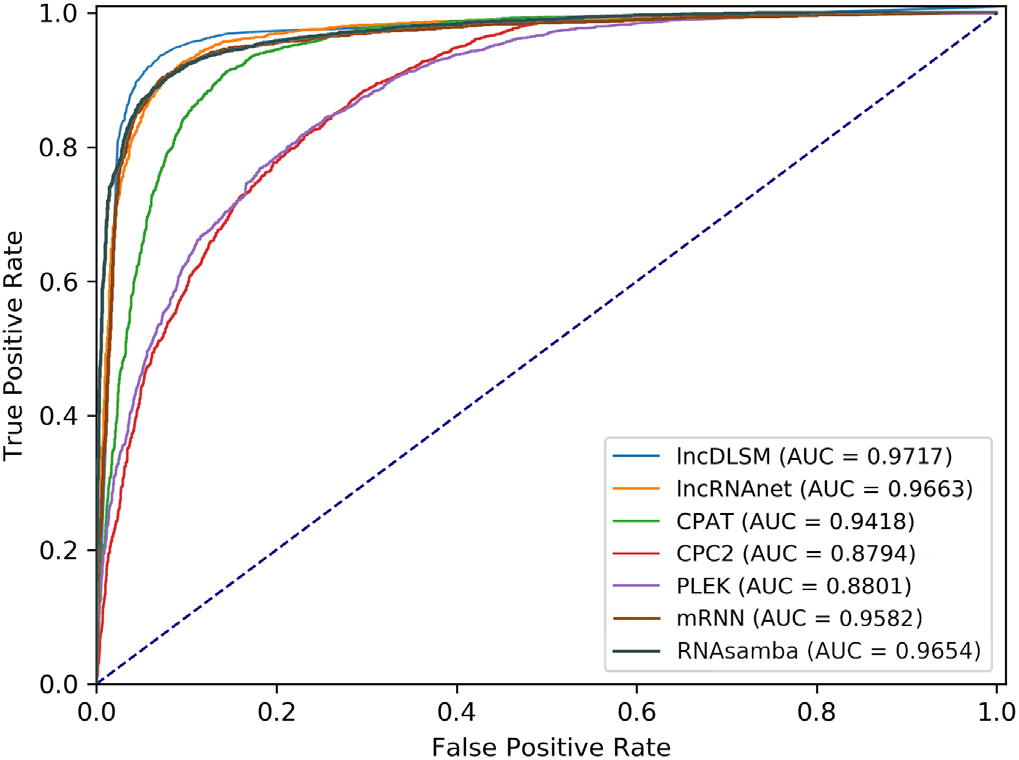
The ROC curves of five different classifiers on the human data

#### 2) Performance comparison of the mouse dataset

**Table V** shows that lncDLSM still achieved good performance with balanced sensitivity and specificity on mouse data. Specially, lncDLSM obtained the highest specificity, accuracy, and F1 and achieved the best overall performance with balanced sensitivity and specificity. Although CPAT had the best performance in sensitivity, its specificity score was not satisfactory. Similarly, CPC2 and PLEK, more inclined to predict lncRNAs, had an imbalanced performance. As shown in **Fig. 4**, 0.9759 AUC score proved that lncDLSM is an excellent classifier.

**Fig. 4.**
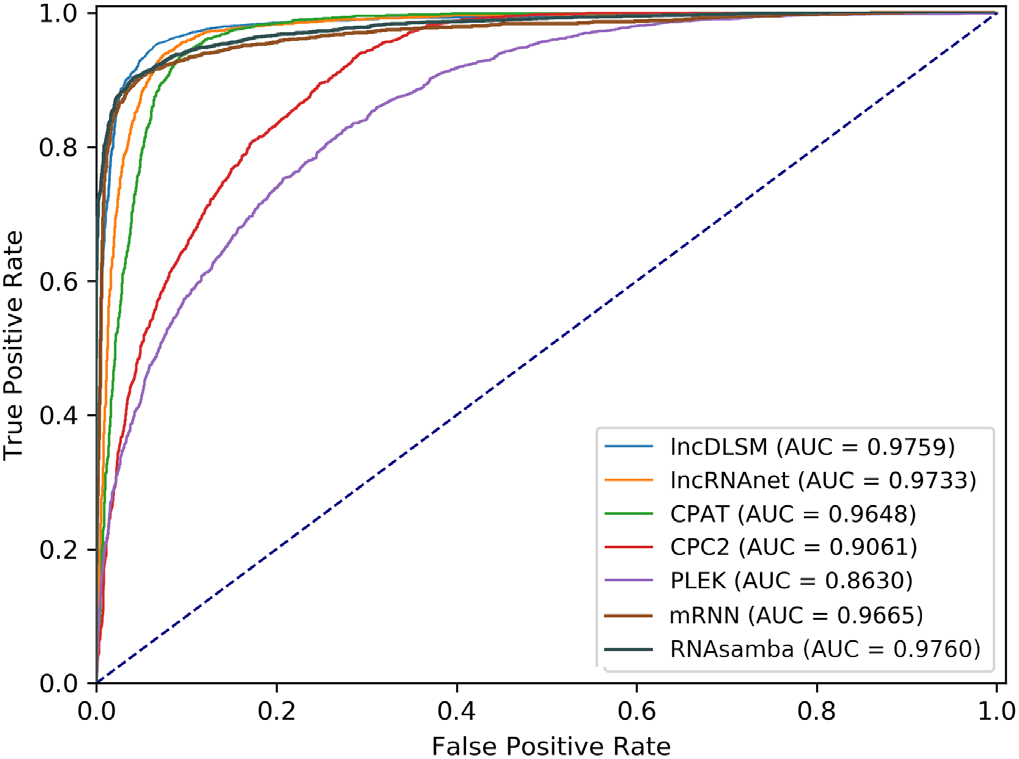
The ROC curves of five different classifiers on the mouse data

**Table V.** Comparison of prediction performance (Species: Mouse)

| Method | Sensitivity | Specificity | Accuracy | F1 |
| --- | --- | --- | --- | --- |
| LncDLSM | 0.9517 | 0.9290 | **0.9403** | **0.9410** |
| LncRNAnet | 0.9623 | 0.8923 | 0.9273 | 0.9298 |
| CPAT | **0.9897** | 0.7753 | 0.8825 | 0.8939 |
| CPC2 | 0.9717 | 0.6493 | 0.8105 | 0.8368 |
| PLEK | 0.9443 | 0.5390 | 0.7417 | 0.7852 |
| mRNN | 0.9050 | **0.9457** | 0.9253 | 0.9238 |
| RNAsamba | 0.9143 | 0.9363 | 0.9253 | 0.9245 |

#### 3) Performance comparison of the other dataset

Additional experimental results with other species are listed in **Table VI**. LncDLSM performed well in sensitivity, specificity, accuracy, F1, and AUC. In particular, lncDLSM achieved more than 94.4% accuracy, 94.4% F1, and 0.976 AUC scores in pig, cow, and rat data. CPAT and CPC2 showed balanced performance on pig, cow, and rat data. In contrast, CPAT and CPC2 performed poorly in human and mouse data. There is a possible reason that the PCTs in the GENCODE have longer 5’ and 3’ Untranslated Regions (UTRs) than those in the RefSeq (Kong, et al., 2007), which contributes to the misjudgment of classifiers. The results demonstrated that lncDLSM and lncRNAnet could ignore the effect of UTRs length, especially lncDLSM. What’s more, **Table V** and **Table VI** reflect that transfer learning can enhance the generalization of lncDLSM.

**Table VI.** Comparison of prediction performance on other species

| Species | Method | Sensitivity | Specificity | Accuracy | F1 | AUC |
| --- | --- | --- | --- | --- | --- | --- |
| Pig | lncDLSM | 0.9483 | **0.9593** | 0.9538 | 0.9536 | 0.9878 |
|  | lncRNAet | 0.9303 | 0.9460 | 0.9382 | 0.9377 | 0.9812 |
|  | CPAT | 0.9637 | 0.9527 | **0.9582** | **0.9584** | 0.9888 |
|  | CPC2 | 0.9517 | 0.9573 | 0.9545 | 0.9544 | **0.9902** |
|  | PLEK | 0.9233 | 0.8577 | 0.8905 | 0.8940 | 0.9592 |
|  | mRNN | 0.9540 | 0.9093 | 0.9317 | 0.9332 | 0.9723 |
|  | RNAsamba | **0.9753** | 0.8980 | 0.9367 | 0.9390 | **0.9902** |
| Cow | lncDLSM | 0.9790 | **0.9853** | **0.9822** | **0.9821** | 0.9962 |
|  | lncRNAet | 0.9890 | 0.9507 | 0.9698 | 0.9704 | 0.9920 |
|  | CPAT | **0.9973** | 0.9420 | 0.9697 | 0.9705 | **0.9980** |
|  | CPC2 | **0.9973** | 0.9463 | 0.9718 | 0.9725 | 0.9978 |
|  | PLEK | 0.9870 | 0.8620 | 0.9245 | 0.9289 | 0.9869 |
|  | mRNN | 0.9413 | 0.9833 | 0.9623 | 0.9615 | 0.9725 |
|  | RNAsamba | 0.9743 | 0.9823 | 0.9783 | 0.9782 | 0.9967 |
| Rat | lncDLSM | 0.9467 | **0.9413** | **0.9440** | **0.9441** | 0.9767 |
|  | lncRNAet | 0.8087 | 0.9297 | 0.8692 | 0.8607 | 0.9481 |
|  | CPAT | 0.9267 | 0.9300 | 0.9283 | 0.9282 | 0.9812 |
|  | CPC2 | 0.9360 | 0.9360 | 0.9360 | 0.9360 | **0.9816** |
|  | PLEK | 0.9357 | 0.7443 | 0.8400 | 0.8540 | 0.9233 |
|  | mRNN | 0.9150 | 0.7490 | 0.8320 | 0.8449 | 0.9158 |
|  | RNAsamba | **0.9647** | 0.7873 | 0.8760 | 0.8861 | 0.9687 |

All the experimental results demonstrated that the transfer learning from the source domain (human data) to the target domain (other species data) was successful, with no negative transfer. It’s conceivable that gene homology among species guarantees successful transfer learning. Indeed, species specificity is the source of nuances in various model parameters. For further explanation, we performed dimensionality reduction on the output of the last hidden layer of neural networks with t-distributed stochastic neighbor embedding (t-SNE) (Kang, et al., 2017). We drew the distributions of lncRNAs/PCTs on five species data. Considering **Fig. 5** and **Fig. 6**, it is easy to see that the small distances and the distinct boundaries among distributions correspond to the homology and the specificity among species, respectively. As we know, human and mouse have high DNA homology. And as shown in Figure 6, the small distance, even a slight overlap between distributions of human and mouse PCTs, is further evidence. Thus, if we can extract features and design the model from the perspective of species homology, transfer learning may be an effective method for revealing relationships among species.

**Fig. 5.**
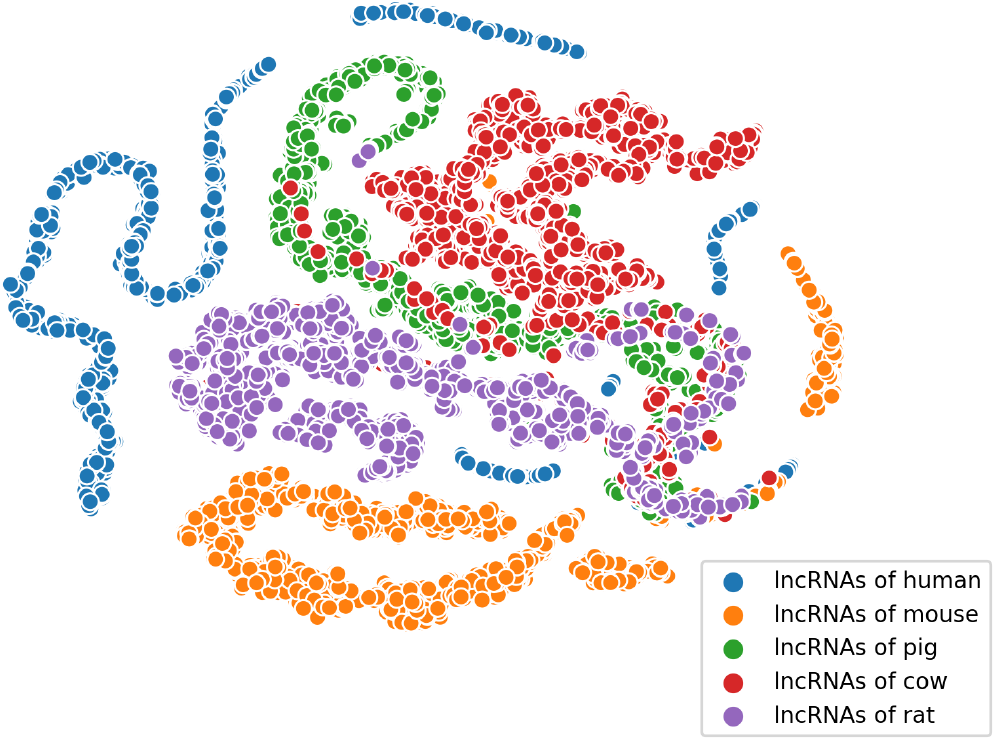
The distributions of lncRNAs in the advanced representation space

**Fig. 6.**
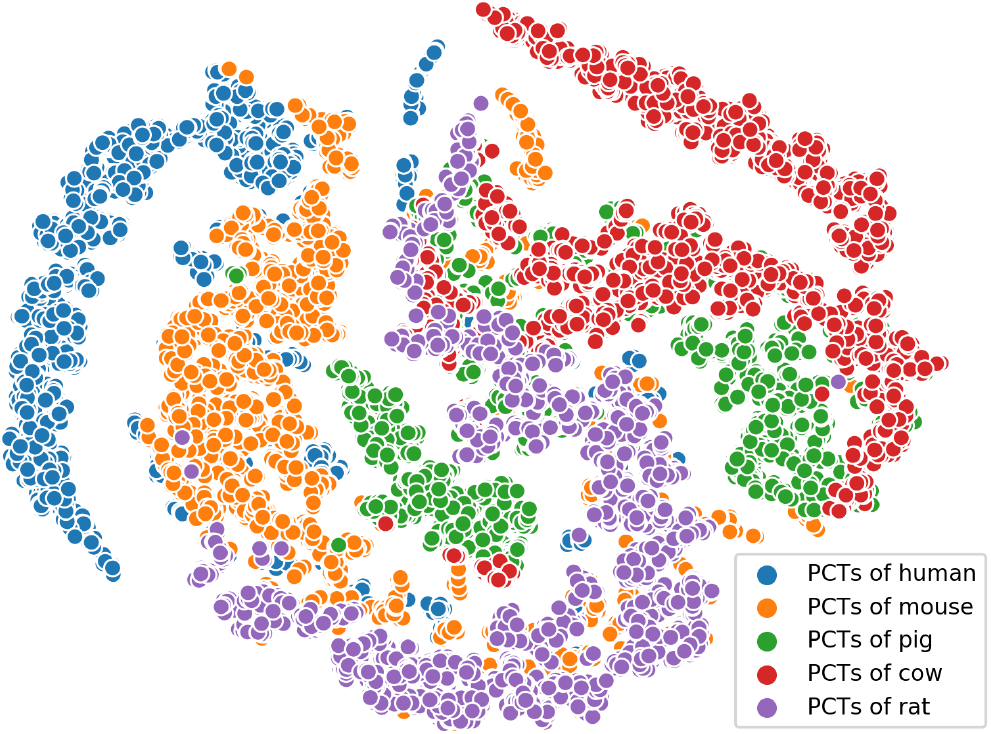
The distributions of PCTs in the advanced representation space

#### C. Online web server

For easy usage of our tool, we also provided an online webserver to predict lncRNAs. We have wrapped up the whole pre-trained models and the detailed algorithms hidden from end users. Input a fasta sequence or upload the candidate sequence into the web and submit the job will quickly give the user prediction results based on the species users choose. End users may check the results online or download the final predictions directly from our web. The web server is freely available at http://39.106.16.168/lncDLSM; our web screenshot is shown in **Fig. 7**.

**Fig. 7.**
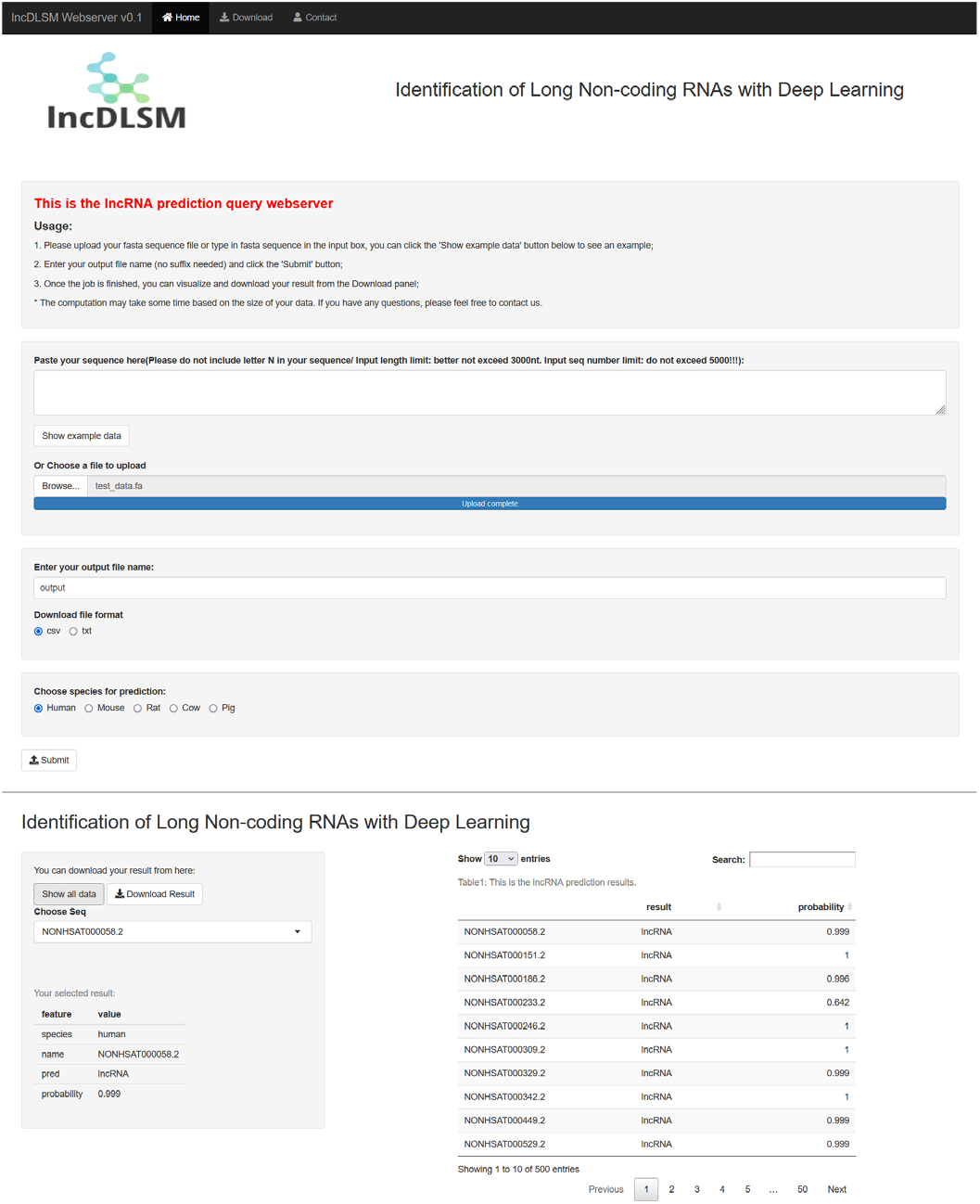
Online lncDLSM webserver

## IV. Discussion

Deep learning has made great strides in computer vision (Y. Li, et al., 2014) and natural language processing (Hill, et al., 2018) with its outstanding characterization ability. In the bioinformatics domain, deep learning-based approaches were also gradually applied (Frankish, et al., 2015; Tsung Yi Lin, et al., 2017; van der Maaten & Hinton, 2008). It is undeniable that prior biological knowledge plays a positive role in identifying lncRNAs to some extent. Still, our deep learning-based approach, lncDLSM, performs well using pure sequence features. Our study provides a new way for sequence characterization.

Prior biological knowledge is empirical knowledge, not a scientific conclusion. In the existing lncRNAs recognition methods, most of the features based on biological knowledge are related to the coding sequences (CDs), which contain one or more ORFs. Thus, these methods focus more on the coding region of sequences and ignore the influences of UTRs. This results in biased predictions toward the lncRNAs. The experimental results show that they perform better when the proportion of CDs region in the whole sequence increases. In contrast, lncRNAnet de-emphasized the use of prior biological knowledge and improved the recognition performance of the model. Specifically, lncDLSM extracted FFT spectrum features and k-mer frequency features to character the sequences’ variations and compositions, respectively, and completely solved the shortcomings of insufficient attention to UTRs. It is worth mentioning that PLEK ignores the sequence variations, resulting in unsatisfactory recognition results.

Even so, lncDLSM has shortcomings, mainly due to the poor interpretability of the deep learning model. In the meantime, in its current form, the biological significance of the advanced features extracted by the model cannot be used. But neural networks combining the transfer learning may be a potential solution to this issue. If the analysis of the features is completed, we can have a more profound understanding of lncRNAs.

## V. Conclusion

With the rapidly intensifying attention on studying long non-coding RNAs, fast and accurate computational methods that can identify lncRNAs from protein coding transcripts remain challenging. In this study, we proposed lncDLSM, a sequence recognition model requiring no prior biological knowledge. CNN-based detector and HINN-based analyzer were designed to mine the variations and composition information of the sequences, respectively. LncDLSM outperformed other state-of-the-art methods such as lncRNAnet, CPAT, CPC2, and PLEK on the HT. For rigor, lncDLSM predicted the other mammals (mouse, pig, cow, and rat) by transfer learning and successfully detected lncRNAs. Specially, all the experimental results showed that lncDLSM has a balanced performance. Thus, lncDLSM is a useful tool for identifying lncRNAs without prior biological knowledge and can be applied to other species by transfer learning. And the success of transfer learning may guide new directions for our further research on the issues related to species homology despite other biological problems. An online web server is also provided for the community for lncRNA prediction and is available for free.

